# Transition bias influences the evolution of antibiotic resistance in *Mycobacterium tuberculosis*

**DOI:** 10.1101/421651

**Authors:** Joshua L. Payne, Fabrizio Menardo, Andrej Trauner, Sonia Borrell, Sebastian M. Gygli, Chloe Loiseau, Sebastien Gagneux, Alex R. Hall

## Abstract

Transition bias, an overabundance of transitions relative to transversions, has been widely reported among studies of mutations spreading under relaxed selection. However, demonstrating the role of transition bias in adaptive evolution remains challenging. We addressed this challenge by analyzing adaptive antibiotic-resistance mutations in the major human pathogen *Mycobacterium tuberculosis*. We found strong evidence for transition bias in two independently curated datasets comprising 152 and 208 antibiotic resistance mutations. This was true at the level of mutational paths (distinct, adaptive DNA sequence changes) and events (individual instances of the adaptive DNA sequence changes), and across different genes and gene promoters conferring resistance to a diversity of antibiotics. It was also true for mutations that do not code for amino acid changes (in gene promoters and the ribosmal gene *rrs*), and for mutations that are synonymous to each other and are therefore likely to have similar fitness effects, suggesting that transition bias can be caused by a bias in mutation supply. These results point to a central role for transition bias in determining which mutations drive adaptive antibiotic resistance evolution in a key pathogen.

**Significance statement:** Whether and how transition bias influences adaptive evolution remain open questions. We studied 296 DNA mutations that confer antibiotic resistance to the human pathogen *Mycobacterium tuberculosis*. We uncovered strong transition bias among these mutations and also among the number of times each mutation has evolved in different strains or geographic locations, demonstrating that transition bias can influence adaptive evolution. For a subset of mutations, we were able to rule out an alternative selection-based hypothesis for this bias, indicating that transition bias can be caused by a biased mutation supply. By revealing this bias among *M. Tuberculosis* resistance mutations, our findings improve our ability to predict the mutational pathways by which pathogens overcome treatment.

## Introduction

Mutation creates genetic variation and therefore influences evolution. Mutation is not an entirely random process, but rather exhibits biases toward particular DNA sequence changes. For example, a bias toward transitions (purine-to-purine or pyrimidine-to-pyrimidine changes), relative to transversions (purine-to-pyrimidine or pyrimidine-to-purine changes), has been widely reported among studies of mutations spreading under relaxed selection (1-5).

Demonstrating the role of such transition bias in adaptive evolution remains challenging, with most existing evidence derived from individual case studies (6-10). Stoltzfus and McCandlish recently reported the first systematic study of transition bias in putatively adaptive evolution, using the repeated occurrence of amino acid replacements in laboratory or natural evolution as evidence that the replacements are adaptive (11). Their meta-analysis provides compelling evidence that transition bias influences adaptive evolution, with transitions observed in at least two-fold excess of the null expectation that they occur once for every two transversions. Yet such analyses have two limitations. First, while the repeated occurrence of an amino acid replacement is highly suggestive of adaptation (12), it is not direct evidence of adaptation. Second, an overabundance of transitions could result from a bias in mutation supply (i.e., mutation-based transition bias), from a greater selective advantage conferred by the amino acid replacements caused by transitions relative to those caused by transversions (i.e., selection-based transition bias), or from both. For example, the spontaneous deamination of methylated cytosines to thymines may contribute to mutation-based transition bias (13), whereas the propensity of non-synonymous transitions to conserve the biochemical properties of amino acids better than non-synonymous transversions may contribute to selection-based transition bias (14). Discriminating amongst these two forms of transition bias has proven challenging to date (15, 16).

Despite the emerging evidence that transitions are overrepresented among adaptive mutations as well as those spreading under relaxed selection, it remains unclear whether this plays a role in real-world scenarios where rapid adaptive evolution has important implications for treatment of infectious disease, such as the emergence of antibiotic resistance in pathogenic bacteria. For example, if pathogen populations fix the first resistance mutation that appears (‘first-come-first-served’) (17), then a mutation supply biased toward particular types of nucleotide substitutions will influence which genetic changes drive adaptation. Alternatively, if many beneficial mutations are available to selection, and pathogens fix those with the highest selective advantage (‘pick-the-winner’), a bias in mutation supply would have a weaker impact on which genetic changes drive adaptation. Therefore, identifying transition bias in such scenarios would improve our basic understanding of how resistance evolves, and our ability to predict the relative likelihoods of alternative mutational pathways to resistance.

Here, we study transition bias in the evolution of antibiotic resistance in *Mycobacterium tuberculosis* (MTB), a major human pathogen for which antibiotic resistance evolution is a key obstacle to effective treatment (18). We do so using two independently curated datasets of mutations that are known to confer antibiotic resistance and are therefore definitively adaptive. Additionally, we test specifically for mutation-based transition bias by considering two subsets of adaptive mutations: 1) Mutations located in gene promoters and in the ribosomal gene *rrs*, which is not translated to protein and therefore should not be influenced by selection-based bias caused by transitions encoding different amino acid changes than transversions. 2) Mutations that are synonymous to each other and are therefore likely to have similar fitness effects.

Our results reveal strong transition bias in the mutational paths to antibiotic resistance and in the number of times each mutational path is used in the evolution of antibiotic resistance, across 22 genes or gene promoters that confer resistance to 11 antibiotics. We also observe transition bias among adaptive mutations that do not code for amino acid changes, and among adaptive mutations that are synonymous to each other, consistent with the hypothesis that transition bias is at least partly mutation-based. We therefore demonstrate that transition bias influences adaptive evolution, specifically the evolution of antibiotic resistance in a key global pathogen.

## Results

We curated a dataset of 152 unique point mutations that confer resistance to at least one of 11 different antibiotics, and that appeared in at least one of 9,351 publicly-available MTB genomes (Materials and Methods). We refer to this dataset as the *Basel dataset*. We also analyzed an independently curated dataset of 208 unique point mutations that confer resistance to at least one of 8 antibiotics and appeared in at least one of 5,310 MTB genomes (19) (Materials and Methods). We refer to this dataset as the *Manson dataset*. The Basel and Manson datasets have 64 point mutations in common, and together include resistance mutations for 11 antibiotics.

Following Stoltzfus and McCandlish (11), we separately studied transition bias in mutational *paths* and in mutational *events*. A mutational path is any single mutation in one of our datasets, such as the well known C > G transversion that causes the S315T substitution in the sole MTB catalase KatG to confer resistance to isoniazid (20). This mutational path may be used any number of times during adaptation of MTB strains to antibiotics, for example in different patients or in different geographic regions. Each use is a mutational event. For the Basel dataset, we calculated the number of mutational events using a parsimony-based analysis of mutational gains and losses at all nodes in the reconstructed phylogeny of the 9,351 MTB genomes (Materials and Methods). The Manson dataset also contained estimates of the number of mutational events, derived using similar methods (Materials and Methods).

We studied transition bias by calculating the ratio of the number of transitions to the number of transversions in each dataset, separately for mutational paths and events. We calculated 95% binomial confidence intervals on this ratio and an empirical *p*-value that describes the probability of observing a transition:transversion ratio greater than the observed ratio, given a null model (Materials and Methods). Among several null models described by Stoltzfus and McCandlish (11), we chose the most conservative, which assumes that all nucleotide mutations are equally likely and there is no difference between the average fitness effects of transitions and transversions. This null model gives a transition:transversion ratio of 0.5, because for any given nucleotide there is one possible transition and two possible transversions. We therefore inferred transition bias in cases where the lower bound of the 95% binomial confidence interval of the observed transition:transversion ratio was greater than 0.5 and the empirical *p*-value was less than 0.05.

We observed transition bias in both mutational paths and mutational events, for both the Basel and Manson datasets (Figure 1). Transition bias was considerably more pronounced in the number of mutational events, as compared to the number of mutational paths. This provides compelling evidence that transition bias influences the evolution of antibiotic resistance in MTB. To understand why, assume there was no mutation-based or selection-based transition bias and the high transition:transversion ratio at the level of mutational paths was instead due to chance, such as a sampling artifact. In this scenario we would expect at most the same transition bias at the level of events, rather than the roughly two-fold increase that we observed.

**Figure 1.**
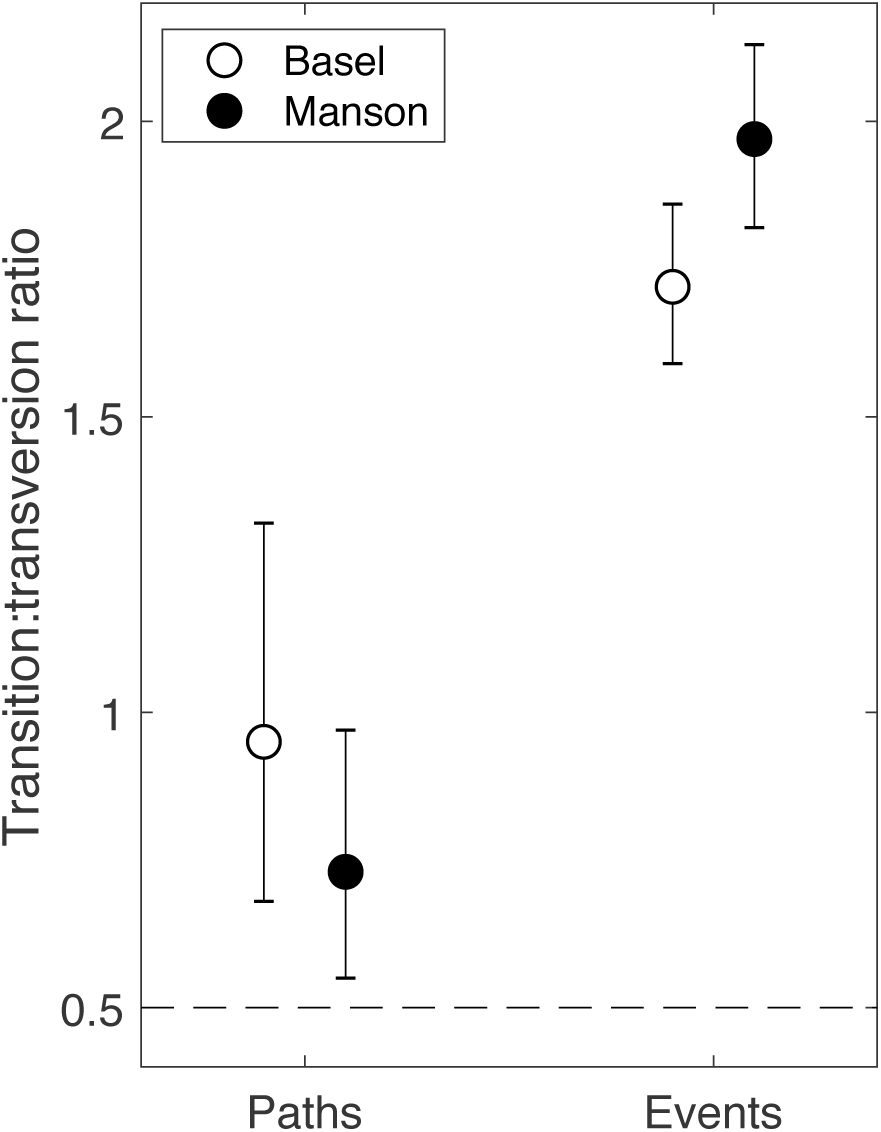
Transition bias in mutational paths and mutational events in the Basel and Manson datasets. The dashed horizontal line shows the null expectation of the transition:transversion ratio.

To determine which mutations explained the observed transition bias, we calculated the relative rates of all six possible nucleotide pair mutations, accounting for GC content (1), which is relatively high in MTB compared to other bacteria (Materials and Methods). For mutational paths, the rate of A/T > G/C transitions exceeded the rate of any other nucleotide-pair mutation (Figure 2A,B). The same was true for mutational events (Figure 2C,D), where the rates of both forms of transitions (G/C > A/T and A/T > G/C) were at least 1.5 times that of all forms of transversions.

**Figure 2.**
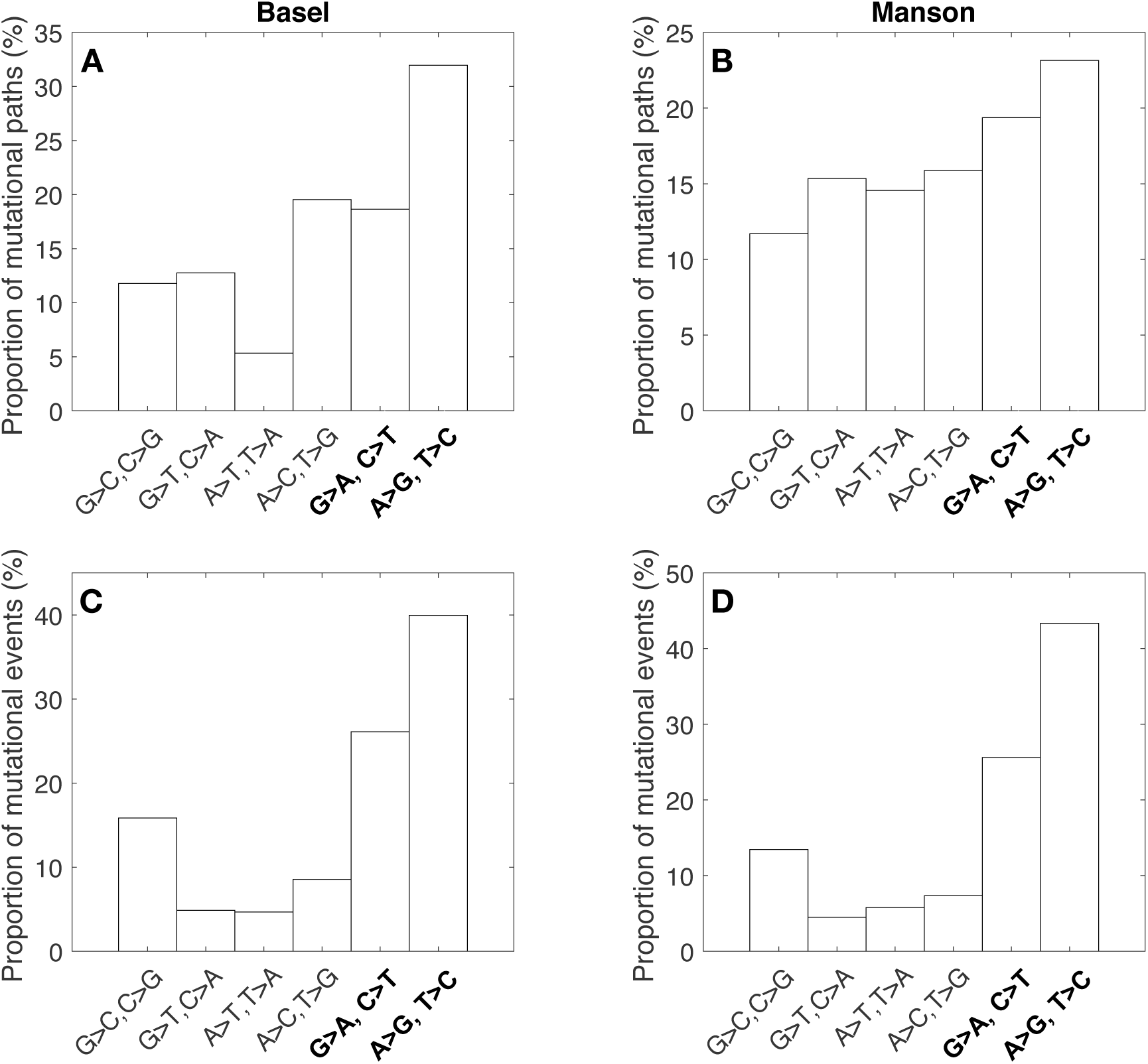
Relative rates of the six nucleotide pair mutations, for mutational paths and events in the Basel and Manson datasets. Transitions are indicated with bold text. Rates adjusted for GC content (Materials and methods).

Because the influence of transition bias might depend on the mechanism of antibiotic resistance, we next tested for transition bias separately for different antibiotics. This reduced the number of mutational paths and events that could be analyzed, so we first determined the antibiotics for which we had sufficient statistical power to ensure that an observed lack of transition bias was not due to reduced sample size (Materials and Methods). This analysis revealed that, at the level of mutational paths, our datasets were too small to test for transition bias of the strength observed in the entire dataset for any individual antibiotic (Figure S1A,B). This is because at least 44 (Basel) and 118 (Manson) mutational paths would be required to provide sufficient statistical power, and the maximum number of paths per antibiotic was 30 (for streptomycin) in the Basel dataset and 51 (for rifampicin) in the Manson dataset (Figure S2A,B). In contrast, we were able to test for transition bias in the mutational events associated with resistance to individual antibiotics, because only 14 and 12 events were required to provide sufficient statistical power in the Basel and Manson datasets (Figure S1C,D). For this analysis, we considered mutations that simultaneously conferred resistance to multiple antibiotics in separate categories. For example, some mutations in the gene Rv1484 (*inhA*) and its promoter confer resistance to both isonazid and ethionamide (5 and 6 mutations respectively in the Basel and Manson datasets). To avoid counting such mutations multiple times, we analyzed them separately and did not include them together with mutations that conferred resistance only to isonazid or ethionamide. All but two individual antibiotics, both in the Basel dataset, had enough mutational events to test for transition bias (Figure S2C,D).

We observed transition bias in the number of mutational events associated with the evolution of resistance to almost all antibiotics, with transition bias indicated by transition:transversion ratios ranging from 1.59 for pyrazinamide to 41.6 for kanamycin (i.e., from more than three-fold to more than eighty-fold excess of the null expectation; Table 1). The glaring exception was isoniazid, which was dominated by a single C > G transversion (S315T in KatG), representing 409 of the 472 mutational events associated with isoniazid resistance in the Basel dataset (or 671 events if we include mutations in *inhA* that are associated with resistance to both isoniazid and ethionamide) and 321 of the 389 mutational events associated with isoniazid resistance in the Manson dataset (or 545 if we include *inhA*).

**Table 1.** Summary of transition bias in mutational events per antibiotic in the Basel and Manson datasets. Rows are ordered by decreasing number of events in the Basel dataset. Dashes indicate antibiotics for which there are no mutational events in the respective dataset. Mutations that confer resistance to multiple antibiotics are reported separately and were not counted among the events conferring resistance to individual antibiotics. RIF-Rifampicin, EMB -Ethambutol, INH – Isonazid, SM – Streptomycin, ETH - Ethionamide, FQ – Floroquinolones, AK – Amikacin, CAP – Capreomycin, KAN – Kanamycin, PZA – Pyrazinamide, OFX - Ofloxacine. T_*i*_ – number of transitions, T_*v*_ – number of transversions, T_*i*_ : T_*v*_ – transition:transversion ratio, 95% CI – 95% binomial confidence interval, *p* – empirical *p*-value.

|  | Basel dataset |  |  |  |  | Manson dataset |  |  |  |  |
| --- | --- | --- | --- | --- | --- | --- | --- | --- | --- | --- |
| Antibiotic | $T_i$ | $T_v$ | $T_i:T_v$ | 95% CI | $p$ | $T_i$ | $T_v$ | $T_i:T_v$ | 95% CI | $p$ |
| RIF | 365 | 223 | 1.64 | (1.38, 1.94) | 0 | 494 | 209 | 2.36 | (2.01, 2.80) | 0 |
| EMB | 390 | 154 | 2.53 | (2.10, 3.07) | 0 | 342 | 131 | 2.61 | (2.13, 3.22) | 0 |
| INH | 42 | 430 | 0.10 | (0.07, 0.13) | 1 | 54 | 335 | 0.16 | (0.12, 0.22) | 1 |
| SM | 285 | 71 | 4.01 | (3.08, 5.28) | 0 | 254 | 72 | 3.53 | (2.71, 4.65) | 0 |
| INH, ETH | 231 | 31 | 7.45 | (5.11, 11.22) | 0 | 137 | 18 | 7.61 | (4.64, 13.23) | 0 |
| FQ | 166 | 57 | 2.91 | (2.14, 4.01) | 0 | - | - | - | - | - |
| AK, CAP, KAN | 128 | 0 | ∞ | (34.20, ∞) | 0 | - | - | - | - | - |
| PZA | 67 | 28 | 2.39 | (1.52, 3.86) | 0 | 46 | 29 | 1.59 | (0.98, 2.61) | 0 |
| KAN | 52 | 9 | 5.78 | (2.82, 13.34) | 0 | 208 | 5 | 41.6 | (17.54, 129.46) | 0 |
| ETH | 7 | 10 | 0.70 | (0.23, 2.04) | 0.3 | 16 | 10 | 1.60 | (0.68, 3.95) | 0.003 |
| KAN, CAP | 17 | 0 | ∞ | (4.13, ∞) | 0 | - | - | - | - | - |
| OFX | - | - | - | - | - | 220 | 91 | 2.42 | (1.89, 3.12) | 0 |

The above results suggest transition bias influences the evolution of antibiotic resistance in MTB. However, it remains unclear whether this bias is mutation-based or selection-based. To disentangle these potential sources of transition bias, we used two different approaches. First, if the observed bias were caused by transitions and transversions encoding amino acid changes with different average fitness effects, we would not expect the bias to extend to mutations that do not encode amino acid changes. We tested this by examining mutations in gene promoters and the ribosomal gene *rrs*. There were 23 such mutations in the Basel dataset, and 25 in the Manson dataset (12 occur in both datasets). Because this did not provide enough statistical power to perform the analysis at the level of paths, we only considered events here. We found an excess of transitions, demonstrating transition bias among mutations that do not encode amino acid changes. Specifically, in the Basel dataset, there were 603 events that comprised 525 transitions and 78 transversions (transition:transversion ratio of 6.73; 95% binomial confidence interval: (5.30, 8.65); empirical *p*-value = 0). In the Manson dataset, there were 520 events that comprised 421 transitions and 99 transversions (transition:transversion ratio of 4.25; 95% binomial confidence interval: (3.41, 5.35); empirical *p*-value = 0). Note that mutations in genes such as *rrs* may nevertheless have variable fitness effects despite not encoding amino acid changes (21), and we discuss this further below.

Second, we considered cases where different mutations caused the same resistance-conferring amino acid change, and were therefore synonymous to each other and expected to have similar fitness effects. Specifically, we considered amino acid changes that can be caused by both transition and transversion mutations at the same ancestral codon. For example, methionine can mutate to isoleucine via the transition ATG > ATA or the transversions ATG > ATT and ATG > ATC. In the standard genetic code, there are five such amino acid changes (Table 2). These were rare or non-existent in the Basel and Manson datasets (Table 2), except the amino acid change methionine-to-isoleucine, which occurred in 3 mutational paths and 137 events in the Basel dataset, and in 4 mutational paths and 135 events in the Manson dataset. In the Basel dataset, the 137 events comprised 88 transitions and 49 transversions (transition:transversion ratio of 1.80; 95% binomial confidence interval: (1.25, 2.60); empirical *p*-value = 0). In the Manson dataset, the 135 events comprised 96 transitions and 39 transversions (transition:transversion ratio of 2.46; 95% binomial confidence interval: (1.68, 3.67); empirical *p*-value = 0). This shows transitions were also overrepresented among events conferring resistance via the same amino acid change (from methionine to isoleucine).

**Table 2.** Observed transitions and transversions in mutational events in the Basel and Manson datasets, and in mutational paths in the TBDReaMDB dataset, among amino acid changes that can be caused by both transition and transversion mutations to the same codon. Abbreviations as in Table 1.

| Amino acid change | Ancestral codon | Possible Ti:Tv | Basel Ti:Tv | Manson Ti:Tv | TBDReaMDB Ti:Tv |
| --- | --- | --- | --- | --- | --- |
| G → R | GGA | 1:1 | 0:0 | 0:0 | 0:1 |
| G → R | GGG | 1:1 | 0:0 | 0:0 | 1:0 |
| M → I | ATG | 1:2 | 88:49 | 96:39 | 6:5 |
| F → L | TTT | 1:2 | 0:0 | 0:0 | 1:0 |
| F → L | TTC | 1:2 | 0:0 | 0:0 | 1:6 |
| W → R | TGG | 1:1 | 0:0 | 0:0 | 7:0 |
| Stop → R | TGA | 1:1 | 0:0 | 0:0 | 0:0 |

Ideally, we would have sufficient mutational paths or events to perform the above analysis on all of the amino acid changes in Table 2. To this end, we analyzed an older dataset, TBDReaMDB (22), that included a greater number of resistance mutations, but was not compiled with the same strict inclusion criteria as the Basel and Manson datasets. Across the 717 mutational paths from this dataset that we included (Materials & Methods), we observed transition bias (transition:transversion ratio of 0.86; 95% binomial confidence interval: (0.74, 1.00); empirical *p*-value = 0). In this dataset there were also a sufficient number of the amino acid changes shown in Table 2 to take an aggregated approach to calculating transition bias amongst mutations that are synonymous to each other. Specifically, amongst all such mutational paths, there were 16 transitions and 12 transversions (Table 2; transition:transversion ratio of 1.33; empirical *p*-value = 0.014). This was more than twice the expected ratio of 0.63, derived from a null model accounting for the number of transitions and transversions in these particular mutational paths (Materials and Methods).

## Discussion

We found strong transition bias at multiple levels (mutational paths and events) and in multiple independently curated datasets (Basel, Manson, and TBDReamDB). By analyzing antibiotic resistance mutations that confer a fitness benefit in the presence of selecting antibiotics, we overcame the difficulty of classifying mutations as adaptive. By focusing part of our analysis on nucleotide substitutions in gene promoters and the ribosomal gene *rrs*, and on mutations that are synonymous to each other, we overcame the difficulty of ruling out an overabundance of transitions caused by transitions and transversions encoding amino acid changes with different average fitness effects. Our data also revealed notable exceptions to the general trend toward transition bias. In particular, the most common resistance mutation by far against isoniazid was a transversion. Our results have four key implications for antibiotic resistance evolution and mutational biases.

First, quantifying the overabundance of transitions improves our ability to predict mutational pathways to resistance. MTB often acquires multiple resistance mutations sequentially, and some mutational trajectories are more common than others (23). Our results suggest the probability of following a given trajectory will be higher when it contains a greater fraction of transitions than alternative trajectories encoding similar resistance phenotypes.

Second, for at least part of our data, we excluded an important potential explanation for transition bias, specifically that transitions encode amino acid changes with different average fitness effects than transversions. We did this by showing that the bias extended to mutations in gene promoters and the ribosomal gene *rrs*, and to mutations that are synonymous to each other. We still cannot rule out variable fitness effects that are not linked to amino acid changes, as observed for streptomycin-resistance mutations in ribosomal genes (21), or synonymous resistance mutations in other bacterial species (24). Nevertheless, if such effects were important drivers of resistance evolution in MTB, we would expect a higher fraction of observed mutations to be synonymous (0 and 2 mutations in coding regions were synonymous in the Basel and Manson datasets, respectively). Moreover, a recent survey of experimentally observed fitness effects did not support a difference in average fitness effects between transitions and transversions (16). By contrast, analysis of mutations under relaxed selection indicates transitions occur more frequently than transversions, including for MTB (1). Therefore, while we do not argue there is no role for biased fitness effects in general (in fact there is evidence of this for viruses (15)), it is unlikely to be the sole cause of the overabundance of transitions we observed, indicating this is explained instead or in addition by a higher mutation supply of transitions than transversions.

Third, if we then accept that a biased mutation supply explains at least some of the observed bias among mutational paths to resistance, this is consistent with a role for mutation-limited ‘first-come-first-served’ dynamics (17) in resistance evolution in MTB. This is not necessarily the case for other pathogens, such as bacteria where resistance is horizontally transmissible. Indeed, this scenario may be relatively likely for pathogens like MTB, where infections expand slowly from a small infectious dose (25) initially containing little genetic variation. On the other hand, genomic evidence shows multiple resistant MTB lineages can occur within the same patient (26, 27), which is less supportive of mutation-limited dynamics. The extent to which such lineages compete with each other will depend on the spatial population structure, for example if they are in different lung sections they may not compete directly (28). Overall, this indicates resistance may evolve via a process somewhere along a continuum between first-come-first-served and pick-the-winner (29).

Fourth, MTB may be closer to one end of this continuum when challenged with some antibiotics than others. Specifically, a single transversion accounted for >70% of the mutational events conferring isoniazid resistance (the same was also true of mutations conferring resistance to both isoniazid and ethionamide), indicating a greatly reduced role for transition bias. Instead, the high frequency of the S315T mutation in *katG* among isoniazid-resistant isolates may reflect that it confers resistance to clinically relevant concentrations, particularly in combination with *inhA* promoter mutations (30), without severely reducing catalytic function (20). MTB strains carrying this mutation, while not detected in *in vitro* mutant screens (31), also appear more likely to transmit than strains with other isoniazid-resistance mutations (32).

Our observation that transition bias is prevalent at the level of paths, but even stronger at the level of events, is consistent with Stoltzfus & McCandlish’s (2017) observations for mutations in parallel experimental or natural populations, and is expected under mutation-based transition bias when the sample size is large (11). Specifically, when the sample size is small, each observed mutational path is represented by one event only, so the transition:transversion ratio of paths and events is identical. With increasing sample size, individual paths are represented by multiple events. In this case, the transition:transversion ratio among events will tend toward that of the adaptive mutation rate, and transitions will be overrepresented if there is mutation-based transition bias. However the transition:transversion ratio of observed paths will tend toward the ratio for all possible unique adaptive mutations, which would approximate 0.5 even when there is mutation-based transition bias (11). Note the observed ratio at the level of paths, while weaker than at the level of events, still exceeded 0.5. This could be because our datasets are incomplete samples of the possible paths to resistance in MTB, and the likelihood of a given path featuring in our datasets is higher if it occurs more frequently (i.e. mutation-based bias and intermediate sample size). Alternatively, our datasets may capture the vast majority of paths to resistance, but due to selection-based bias a greater fraction of them are transitions than transversions.

The bias we observed was also prevalent at the level of individual nucleotide changes, with A/T > G/C transitions being particularly common. This is in contrast to earlier evidence that G/C > A/T transitions are the most common type of mutation spreading under relaxed selection in multiple species, including MTB (1). Nevertheless, in that study the ratio of transitions to A/T compared to G/C was lower for MTB than other bacteria. Further evidence of a distinctive transition bias toward G/C in Mycobacteria, and in particular an overabundance of A/T > G/C transitions, comes from a recent mutation accumulation experiment with the related species *Mycobacterium smegmatis* (33). Another earlier study testing for evidence that oxidative damage caused by antibiotics influences mutational biases also found that A/T > G/C transitions were the most common type of substitution in an earlier version of the TBDReaMDB (34). Thus, transition bias is pervasive across earlier studies and our dataset, yet may be toward different types of transitions depending on the species and on whether mutations are involved in antibiotic resistance.

In conclusion, our data support the hypothesis that a bias toward transitions plays a key role in determining the genetic changes driving antibiotic resistance evolution.

## Materials and Methods

### The Basel dataset

We curated a list of mutations known to confer resistance to one or more of the following drugs or drug classes: isoniazid, ethionamide, rifampicin, ethambutol, pyrazinamide, fluoroquinolones, aminoglycosides. Starting from a previously published set of 120 mutations (35), we first excluded mutations in *rpsA* and *ahpC*, because these genes were unlikely to confer resistance to pyrazinamide and isoniazid respectively (36, 37). We then added *gyrB* to the list of pertinent genes, because some mutations in this gene have been shown to lead to fluoroquinolone resistance (38).

We included additional mutations if they met one or more of the following criteria: 1) they have been shown, by virtue of allelic exchange, to confer resistance; 2) introduction of the mutation into the enzyme of interest was investigated *in vitro* and shown to confer properties consistent with drug resistance; 3) they were identified in laboratory-generated antibiotic-resistant strains as the most likely candidate for resistance; or 4) there was a clear correlation between the presence of the mutation and drug resistance as detected by phenotypic drug susceptibility testing of clinical strains. This resulted in a list of 196 mutations (Table S1), of which we found 152 in at least one of 9,351 publicly-available MTB genomes (Table S2) (39). These are the mutational paths in the Basel dataset (Table S3).

We determined the mutational events by first calling single nucleotide polymorphisms in the 9,351 genomes and then using the polymorphisms to reconstruct the genomes’ phylogeny and to infer mutational gains and losses throughout the phylogeny, as follows. We clipped Illumina adaptors, trimmed low quality reads using Trimmomatic v. 0.33 (SLIDINGWINDOW:5:20) (40), and removed reads shorter than 20 base pair. We merged overlapping paired-end reads using SeqPrep v. 1.2 (overlap size = 15), and mapped the resulting reads to the reconstructed ancestral sequence of the *M. tuberculosis* complex (41) using the mem algorithm of BWA v 0.7.13 (ref (42)). We marked duplicated reads using the MarkDuplicates module of Picard v. 2.9.1, performed local realignment of reads around indels using the RealignerTargetCreator and IndelRealigner modules of GATK v. 3.4.0 (ref (43)), and excluded reads with an alignment score lower than (0.93 × read_length) – (read_length × 4 × 0.07), which corresponds to more than 7 mismatches per 100 base pairs. We called single nucleotide polymorphisms using Samtools v. 1.2 mpileup (44) and VarScan v. 2.4.1 (ref (45)), with the following thresholds: minimum mapping quality of 20, minimum base quality at a position of 20, minimum read depth at a position of 7, minimum percentage of reads supporting the call 90%, maximum strand bias for a position 90%. Genomes were excluded if 1) they had an average coverage < 20x, 2) more than 50% of their single nucleotide polymorphisms were excluded due to the strand bias filter, 3) more than 50% of their single nucleotide polymorphisms had a percentage of reads supporting the call between 10% and 90%, or 4) they contained single nucleotide polymorphisms that belong to different MTB lineages, because this indicates that a mix of genomes was sequenced. Finally, we excluded all genomic positions with more than 10% missing data. This resulted in a final dataset of 300,583 polymorphic positions.

We inferred a phylogenetic tree based on the 300,583 single nucleotide polymorphisms with FastTree (46), using double digit precision and the options -nocat and -nosupport. We extracted the bases at each of the 152 genomic positions in our list of resistance mutations from the vcf file for the 9,351 genomes and assembled them in phylip format. To determine the number of mutational events per mutational path, we reconstructed the nucleotide changes at the 152 genomic positions on the phylogenetic tree rooted with the inferred ancestral sequence of the MTB complex (41). To do this we used the maximum parsimony ACCTRAN algorithm (47) implemented in PAUP* v. 4.0a (ref (47)), giving equal weight to all characters and considering them as unordered.

### The Manson dataset

Manson et al. (19) compiled a list of polymorphisms associated with resistance to eight antibiotics (Supplementary Table 4 in ref (19)), and searched for these polymorphisms in 5,310 MTB genomes. They found 392 of these polymorphisms in at least one genome (Supplementary Table 5 in ref (19)). We filtered these polymorphisms to only include point mutations, resulting in a dataset of 208 mutational paths. Manson et al. (19) calculated the number of events per mutational path by reconstructing the phylogeny of the 5,310 MTB genomes and using a parsimony-based analysis to determine mutational gains and losses throughout the phylogeny. We used their estimates of the number of events per mutational path, as they were reported in Supplementary Table 5 of ref (19).

### The TBDReaMDB dataset

TBDReaMDB is a dataset of 1,178 mutational paths associated with resistance to at least one of 9 antibiotics. We filtered this dataset to only include mutational paths that (1) are non-synonymous point mutations, with both both the ancestral and derived codons reported (709 paths), or (2) point mutations in promoters that are upstream of a gene’s transcription start site (8 paths), and (3) are non-redundant, where we considered two mutational paths redundant if they were the same mutational path and associated with the same gene ID, drug, and codon position (or nucleotide position in the case of promoters). The filtered dataset contains 717 mutational paths.

We used this dataset, which includes a greater number of mutations than the Basel and Manson datasets, but with less strict inclusion criteria, to study transition bias amongst amino acid changes that can be caused by mutational paths that are transitions or transversions, and that arise from the same codon (Table 2). For this purpose, we developed a null model accounting for the number of transitions and transversions in the available mutational paths that cause such amino acid changes. Specifically, for a given combination *i* of amino acid change and ancestral codon, there are *n*_*i*_ mutational paths that are transitions and *p*_*i*_ mutational paths that are transversions. In the TBDReaMDB dataset, there are *m* = 6 such combinations of amino acid changes and ancestral codons (top six rows of Table 2). Some are observed multiple times (at different loci), giving a total of 28 such observations (i.e., the sum of the values in the rightmost column of Table 2). The expected probability of a transition under the null model is 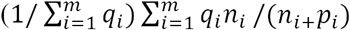, where *q*_*i*_ is the number of times each mutation (path) is observed at different loci (events). For the *m* = 6 observed mutational paths, this gives a null transition probability of (1/(1+1+11+1+7+7))*(1*0.5 + 1*0.5 + 11*0.33 + 1*0.33 + 7*0.33 + 7*0.5) = 0.385, which equates to a transition:transversion ratio of 0.63.

### Calculating empirical *p*-values for transition:transversion ratios

To calculate an empirical *p*-value for an observed transition:transversion ratio in a dataset containing *x* mutational paths or events, we randomly generated 10^6^ datasets of *x* mutations according to our null model that transversions are twice as likely as transitions. Each of the *x* mutations in each dataset was chosen to be a transition with probability 1/3 or a transversion with probability 2/3 (when we analyzed mutations that are synonymous to each other in the TBDReaMDB dataset, we used probabilities of 0.385 and 0.615). We determined the transition:transversion ratio for each of these datasets and then calculated the empirical *p*-value as the fraction of these ratios that were greater than the observed ratio.

### Calculating the relative rates of the six nuleotide pair mutations

GC content may influence the number of mutational paths or mutational events in our datasets. MTB has a high GC content, 65.6% genome-wide (48) and 64.4% in the 17 genes associated with resistance in the Basel and Manson datasets. Thus, we expected to see more mutations from G/C or C/G than from A/T or T/A, simply because there are more Gs and Cs in the genes associated with resistance in our datasets. To control for this effect in our calculation of the relative rates of the six possible nucleotide pair mutations, we followed the method of Hershberg and Petrov (1). Specifically, we first determined the number of mutations of each type we would expect under equal GC content by multiplying the number of mutations from A/T to G/C, C/G, or T/A by 64.4/(100 - 64.4). We then calculated the relative rates of the six nucleotide pair mutations by dividing the number of each (unchanged for mutations from G/C) by the sum of all possible pairs of mutations, and multiplying by 100.

### Calculating statistical power

The numbers of paths and events are much smaller for individual antibiotics than in the full datasets, which may render our test of transition bias statistically underpowered for individual antibiotics. To determine the minumum number of mutational paths or events required to rule out the possibility that an observed lack of transition bias might be due to small sample size, we downsampled our datasets as follows. For each of the Basel and Manson datasets separately, we randomly sampled *x* mutational paths or events from the dataset without replacement and calculated the fraction of these paths or events that were transitions (we calculate this fraction, rather than the transition:transversion ratio, to avoid division by zero) and the associated 95% binomial confidence interval. We repeated this 10^4^ times for each value of *x*, and calculated the average fraction of transitions, the minimum of the lower 95% confidence intervals, and the maximum of the upper 95% confidence intervals. For mutational paths, we varied *x* from 2 to the maximum number of mutational paths in each dataset, and for mutational events, we varied *x* from 2 to 800. We then determined the minimum value of *x* for which the minimum of the 95% confidence intervals exceeded 1/3, which is the expected fraction of transitions under the null model (corresponding to a transition:transversion ratio of 0.5). This is the minimum number of mutational paths or events required to exclude the possibility that an observed lack of transition bias at a given value of *x* is due to small sample size, given the transition bias in the full dataset. For the Basel dataset, this minimum number of mutational paths was 44 and the minimum number of mutational events was 14. In the Manson dataset, the minimum number of mutational paths was 118 and the minimum number of mutational events was 12. No antibiotic was associated with more than these minimum numbers of mutational paths in either the Basel or Manson datasets (Figure S2A,B). However, we had at least the minimum number of mutational events for all antibiotics and combinations except for Amikacin+Capreomycin, and mutations conferring resistance only to Capreomycin in the Basel dataset (Figure S2C,D).

## Acknowledgements

We acknowledge support from Swiss National Science Foundation (SNSF) grants PP00P3_170604 (J.L.P), 31003A_165803 (A.R.H.), 310030_166687 (S.G.), IZRJZ3_164171 (S.G.), IZLSZ3_170834 (S.G.) and CRSII5_177163 (S.G.), the European Research Council (309540-EVODRTB to S.G.) and SystemsX.ch. Part of the calculations were performed at sciCORE (http://scicore.unibas.ch/) scientific computing center at University of Basel.

---

**Figure S1.**
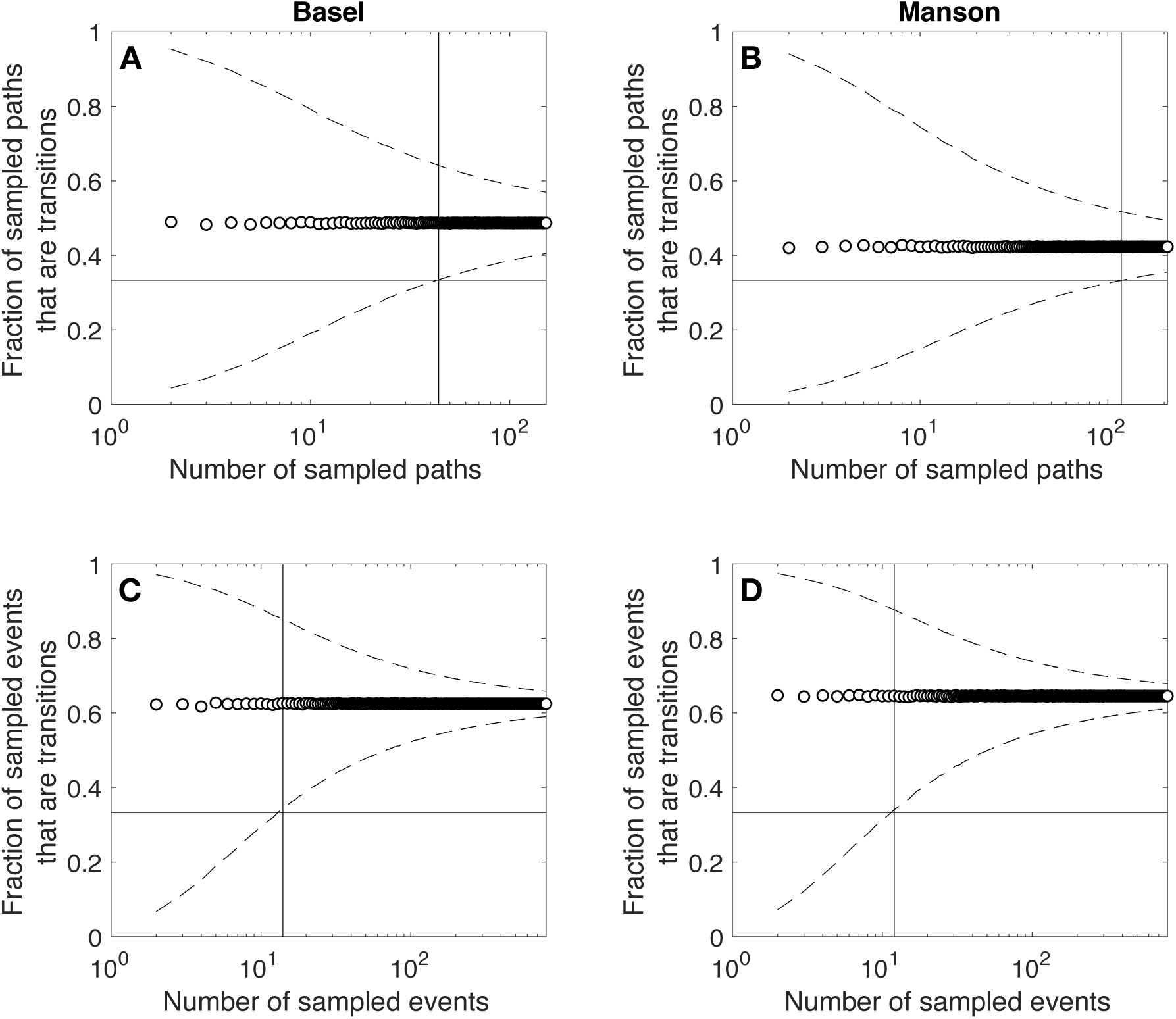
The average fraction of transitions (open circles), with the minimum and maximum of the lower and upper 95% binomial confidence intervals (dashed lines), relative to the number of sampled paths or events from the Basel or Manson datasets (x-axis). Open circles and dashed lines are derived from 10^4^ replications per number of sampled paths or events. The horizontal line indicates the null expectation of a fraction of transitions equal to 1/3, and the vertical line indicates the minimum number of mutational paths or events required for the minimum lower bound on the 95% confidence interval to exceed 1/3.

**Figure S2.**
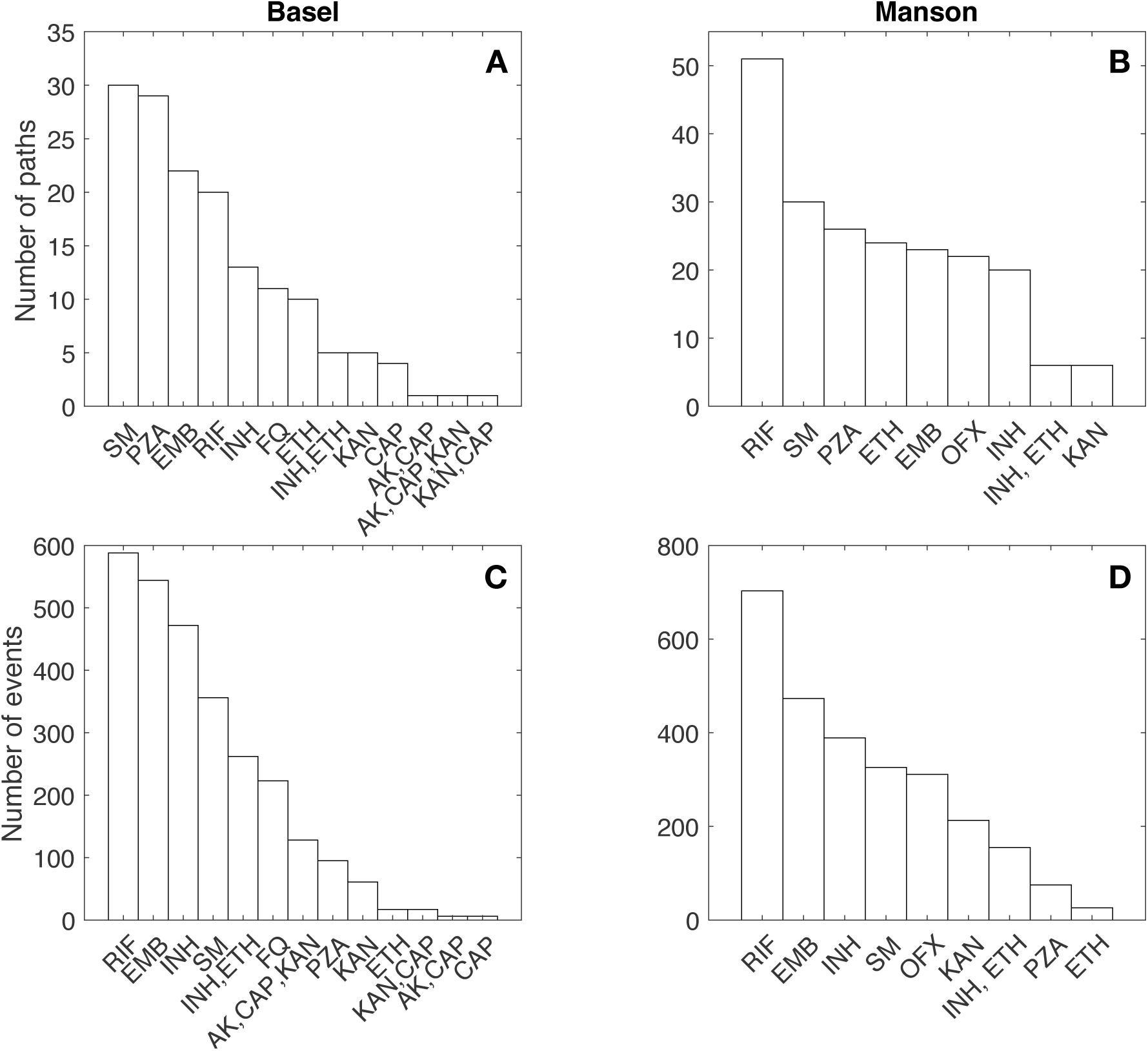
The numbers of mutational paths and events that confer resistance to individual or multiple antibiotics in the Basel and Manson datasets.

